# Optimizing systemic insecticide use to improve malaria control

**DOI:** 10.1101/621391

**Authors:** Hannah R. Meredith, Luis Furuya-Kanamori, Laith Yakob

**Affiliations:** Department of Disease Control, Faculty of Infectious and Tropical Diseases, London School of Hygiene and Tropical Medicine, London, United Kingdom; Research School of Population Health, College of Health and Medicine, Australian National University, Canberra, Australia

**Keywords:** systemic insecticide, malaria, mosquito, vector control, computational modelling

## Abstract

Long lasting insecticidal nets and indoor residual sprays have significantly reduced the burden of malaria. However, several hurdles remain before elimination can be achieved: mosquito vectors have developed resistance to public health insecticides, including pyrethroids, and have altered their biting behaviour to avoid these indoor control tools. Systemic insecticides, drugs applied directly to blood-hosts to kill mosquitoes that take a blood meal, offer a promising vector control option. To date, most studies focus on repurposing ivermectin, a drug used extensively to treat river blindness. There is concern that over-dependence on a single drug will inevitably repeat past experiences with the rapid spread of pyrethroid resistance in malaria vectors. Diversifying the arsenal of systemic insecticides used for mass drug administration would improve this strategy’s sustainability. Here, a review was conducted to identify systemic insecticide candidates and consolidate their pharmacokinetic/pharmacodynamic properties. The impact of alternative integrated vector control options and different dosing regimens on malaria transmission reduction are illustrated through a mathematical model simulation. The review identified drugs from four classes commonly used in livestock and companion animals: avermectics, milbemycins, isoxazolines, and spinosyns. Simulations predicted that isoxazoline and spinosyn drugs were promising candidates for mass drug administration, as they were predicted to need less frequent application than avermectins and milbemycins to maintain mosquitocidal blood concentrations. These findings will provide a guide for investigating and applying different systemic insecticides to achieve better mosquito control strategies.

**Significance:** The widespread use of long lasting insecticidal nets (LLINs) and indoor residual spray has selected for mosquitoes that are resistant to pyrethroids or avoid exposure by feeding outdoors or on livestock. Systemic insecticides, drugs that render a host’s blood toxic to feeding mosquitoes, could be an effective control strategy for mosquitoes with pyrethroid resistance and/or outdoor feeding tendencies. Here, a number of existing systemic insecticide candidates are identified and their pharmacokinetic properties in different drug-host-route scenarios consolidated. These data were used to parameterise a mathematical model that illustrated the projected gains achievable in malaria control programmes already employing LLINs. The findings provide a guide for investigating and applying different systemic insecticides to improve mosquito control strategies and reduce malaria transmission.

## Introduction

Long lasting insecticidal nets (LLINs) and indoor residual sprays (IRS) have played significant roles in reducing the burden of malaria (1, 2). However, several hurdles remain before elimination can be achieved. First, pyrethroids are heavily used in LLINs and, previously, IRS (3). As a result, the widespread and sustained use of this single class of insecticides has selected for mosquitoes that are resistant to the primary intervention methods (4, 5). Second, because LLINs and IRS target mosquitoes that feed indoors on humans, mosquitoes have shifted their feeding patterns to avoid exposure. For instance, increasing numbers of mosquitoes have been found to seek their bloodmeals and/or rest outdoors after a new instalment of bed nets (6). Some malaria-transmitting mosquitoes avoid indoor interventions by obtaining their blood meals from animal hosts (7). Though livestock cannot act as parasite reservoirs, bites diverted away from human hosts can act as temporary reprieve from insecticide exposure, increasing vector lifespans, and consequently contributing to perpetuated transmission (8).

To build on recent gains in malaria vector control, it is critical to develop a method that is effective against pyrethroid resistant, outdoor feeding/resting, or zoophagic mosquitoes (9). A promising solution is systemic insecticides, drugs that render host blood toxic to a mosquito that takes a blood meal (10). The types of systemic insecticides most relevant to treating mosquitoes are ectoparasiticides, drugs that target ectoparasites (e.g. blood-feeding arthropods), and endectocides, drugs that target both endo- and ectoparasites. Many of these drugs have a mode of action distinct from pyrethroids (11–15), and thus should be effective against mosquitoes that either have mutations specific to pyrethroids or are susceptible. These systemic insecticides have been widely used to treat humans, livestock, and domestic animals for infections ranging from gastrointestinal and systemic nematodes to blood-feeding parasites (16–19).

Studies have demonstrated that mass drug administration (MDA) of the systemic insecticide ivermectin to humans and cattle can significantly decrease mosquito population numbers temporarily (20, 21). To attain longer lasting impacts, the optimal use of systemic insecticides requires understanding the pharmacokinetics of the drug in the host to determine the dosing frequency necessary for maintaining lethal blood-drug concentrations. Additionally, understanding the mosquito population’s feeding patterns will guide the decision of whether humans or animal hosts should be targeted. Finally, recent history has highlighted the importance of avoiding over-reliance on singular control tools; thus, it would be prudent to both investigate the effects of synergizing the systemic insecticide with extant interventions and expand the arsenal of effective systemic insecticides from the current candidate of interest, ivermectin (22, 23). Here, a review of endectocides and ectoparasiticides is presented to collate the pharmacokinetic properties for different drug-host-route combinations. These data were then used to parameterise a mathematical model to illustrate the projected gains achievable in malaria control programmes already employing LLINs.

## Methods

### I. Literature review

Veterinary and human parasiticides were identified with systemic properties that affected arthropods and were not prohibited nor being phased out of use in most countries. These included avermectins, milbemycins, neonicotinoids, spinosyns, and isoxazolines. Unlike pyrethroids, which target voltage-gated sodium channels (11), avermectins and milbemycins target glutamate-gated chloride channels (12), neonicotinoids and spinosyns target unique sites of the nicotinic acetylcholinesterase receptor (nAChR) (13, 14), and isoxazolines target γ-aminobutyric acid (GABA)-gated chloride channels(15). Further discussion of the parasiticides’ structures and modes of action is beyond the scope of this study, but may be reviewed elsewhere (12–15).

To determine the relevant pharmacokinetic studies for different systemic insecticide treatments, a review of the electronic literature was conducted. PubMed was searched from inception to July 24, 2018 using the following search terms: (*“systemic insecticide”* OR *endectocide* OR *avermectin* OR *abamectin* OR *doramectin* OR *eprinomectin* OR *ivermectin* OR *selamectin* OR *milbemycin* OR *“milbemycin oxime"* OR *moxidectin, ectoparasiticides* OR *isoxazolines* OR *afoxolaner* OR *fluralaner* OR *sarolaner* OR *lotilaner* OR *neonicotinoids* OR *imidacloprid* OR *nitenpyram* OR *spinosyns* OR *spinosad* OR s*pinetoram* OR "*N-tert-butyl nodulisporamide*") AND (*pharmacokinetics* OR "*area under the curve*" OR "*area under curve*" OR *kinetics* OR "*half-life*" OR *Cmax* OR *Tmax* OR *blood* OR OR *plasma)*. The complete list of accepted and rejected studies is available upon request to the corresponding author.

Study inclusion was determined in two steps. First, the titles and abstracts were screened to determine studies that were not relevant, not primary research (e.g., letter or review), or purely computational. Irrelevant studies were defined as those focusing on drug mechanism, a drug that was not systemic or ineffective against mosquitoes, or hosts that are not targeted by mosquitoes for blood meals. After the initial screen, the full papers of the remaining studies were reviewed. Inclusion required the reporting of relevant pharmacokinetic parameters in plasma, use of a standardized drug (i.e. generic or a commercially available formula), application of single drug (i.e. no adjuvants or cocktails), and a test-population size n ≥ 3.

From the selected studies, the following data were extracted directly into an Excel spreadsheet: host studied, drug applied, drug name/formula, dose applied, route of administration, the maximum drug concentration reached in the plasma (C_max_), the time it took to reach C_max_, the area under the curve, the half-lives for absorption and elimination, the mean residual time, and the volume of distribution. All data were summarized using basic descriptive statistics (mean and standard deviation or standard error when available) for each scenario (host-drug-route of administration). As different classes of drugs require different concentrations to achieve the same toxicity level, the doses for each combination of drug-host-route were not compared. Instead, the analysis focused on the pharmacokinetic metrics that impact the treatment’s efficacy, which in turn, can be used to calculate the appropriate dose.

### II. Data analysis

To determine the underlying trends between the pharmacokinetic parameters and each categorical factor (host, drug class, route of administration), a weighted (using the inverse of the variance) three-way ANOVA was conducted in Stata. Hosts with fewer than 10 observations were grouped into two categories based on bodyweight: other-small (< 75 kg) and other-large (> 75 kg).

### III. Model development

To investigate strategies for applying different systemic insecticides to further limit malaria transmission, models from the literature were modified (24–26) (see **Supplementary Text** for model development and **Table S1** for parameter values). The proportion of bites that lead to infection, egg laying rate, and death rate of mosquitoes are dependent on the concentration of insecticide in LLINs (N) and systemic insecticides in livestock or humans (D_L_ or D_H_, respectively) a mosquito is exposed to. Different host species and routes of administration are characterised by different rates of adsorption, which were often not reported. Hence, to allow for comparison, the model represented the systemic insecticide’s pharmacokinetics as a single compartment and the initial insecticide concentration as the reported C_max_ in the blood after treatment.

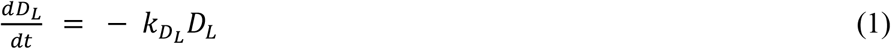

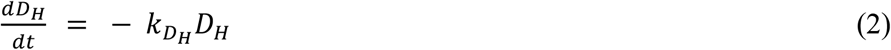

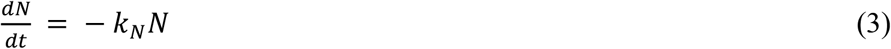

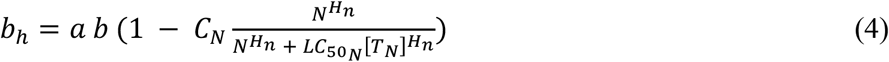

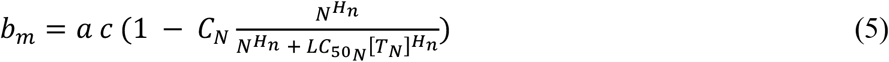

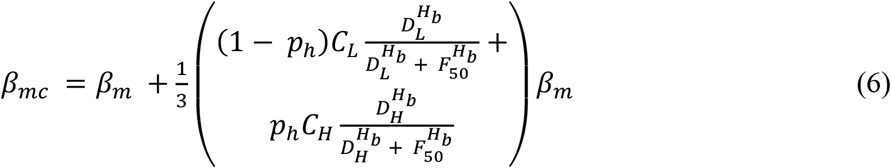

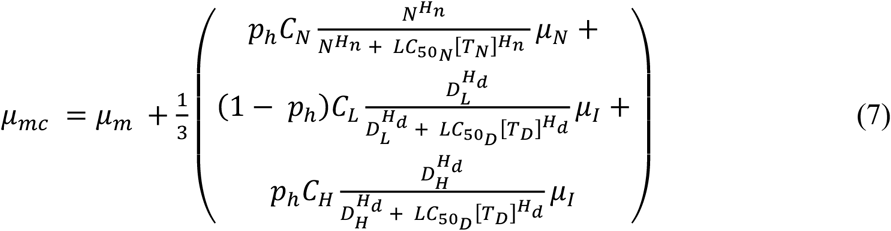

The impact of insecticides used for livestock treatment, human treatment, and bed net treatment is diminished as they degrade at rates 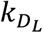, 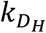, and *k*_*N*_. These rates were determined by the half-lives recorded from the review. The transmission potential from vector to human or vice versa (*b*_*h*_ and *b*_*m*_, respectively) is a function of biting frequency (*a*), the proportion of bites that successfully leads to infection in humans or mosquitoes (*b* and *c*, respectively), the coverage of bed nets (*C*_*N*_) and whether the LLIN’s insecticidal concentration is above the lethal concentration for killing 50% of the population in a set amount of time 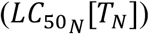. Mosquitoes lay eggs at a natural rate of *β*_*m*_, but the number of eggs laid can be modified with the introduction of certain systemic insecticides (*β*_*mc*_). Similarly, mosquitoes natural death rate (*μ*_*m*_) is modified with the introduction of LLINs and systemic insecticides 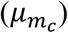. For both the egg laying rate and death rate, the impact of different control strategies depends on the proportion of bites on humans (*P*_*H*_), the coverage of LLINs and systemic insecticides in livestock or humans (*C*_*N*_, *C*_*L*_, *C*_*H*_, respectively), and the respective concentration thresholds for reducing fecundity or killing by 50% (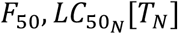, and 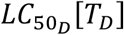).

The relationship between mosquito fecundity or mortality and the concentration of LLINs or systemic insecticides is not linear, but is captured by Hill kinetics. The *F*_50_, 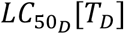, and Hill coefficients (*H*_*b*_ and *H*_*d*_ for birth and death rates, respectively) were calculated for lab-reared *Anopheles* for different systemic insecticides by fitting a Hill equation to published data (27–29) (**Fig. S1-2**). The 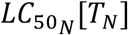 was calculated for wild *Anopheles* that displayed a range of resistance to LLINs and the Hill coefficient (*H*_*n*_) was calculated for lab-reared, permethrin resistant *Anopheles* by fitting a Hill equation to published data (30, 31). The time window (*T*_*N*_ or *T*_*D*_) associated with each insecticide’s *LC*_50_was based on previously reported measurements (27–29, 31).

### IV. Simulations

This model was used to explore the impact of synergizing different systemic insecticide treatments and LLINs on permethrin resistant mosquitoes. A new LLIN was replaced every three years, as recommended by the WHO (32). Systemic insecticide treatments were designed to test a range of half-lives (0.1:100 days), dose concentrations (1:10^5^ ng/mL), F_50_s (0:480 ng/mL), and 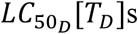 (7:1180 ng/mL), based on data mined from the review. The simulation begins with the initial concentration of drug in the host’s blood, approximated by C_max_. The appropriate dose necessary to achieve C_max_ can be calculated based on pharmacokinetics associated with each host species, as previously documented (33), and is not discussed here. Although many of the systemic insecticides identified in the review remain to be characterised as mosquitocidal candidates, 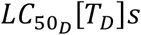 were identified from the avermectin, moxidectin, isoxazoline and spinosyn classes for *Anopheles gambiae* or *An. arabiensis* and used to establish a range of realistic values. *F*_50_*s* were only reported for a subset of avermectins and moxidectin. Each treatment characterised by half-life, C_max_, *F*_50_, and 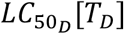 was dosed at frequencies ranging from weekly to annually. Strategy outcome was quantified by calculating the relative reduction (RR) in malaria prevalence after 3 years of drug treatment and LLIN coverage (*M*_*N*+*D*_) relative to 3 years of LLINs alone (*M*_*N*_).

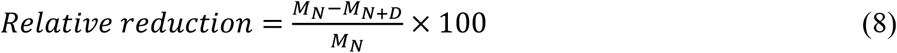

This framework was applied to evaluate strategies for areas with different levels of baseline malaria, mosquito populations with different feeding behaviours, and different deployment scenarios (targeting livestock only, humans only, or both).

## Results

### I. Data retrieval

From the initial 375 articles returned by the search, 237 full-text articles were assessed for eligibility, and 139 met the eligibility criteria (Fig. 1a). The studies reported pharmacokinetic parameters in eight different host categories, ten systemic insecticides, and six routes of administration (Fig. 1b-d). The three most commonly studied hosts were cattle, sheep, and dogs. Systemic insecticides studied included five avermectins (abamectin, doramectin, eprinomectin, ivermectin, and selamectin), three isoxazolines (afoxalaner, fluralaner, and lotilaner), one milbemycin (moxidectin), and one spinosyn (spinosad). These insecticides were applied intramuscularly, intraruminally, intravenously, orally, subcutaneously, or topically. Note that intraruminal and intravenous routes of administration are experimental and are not currently operationally feasible; however, they were included to help determine the full range of action possible for each drug.

**Fig. 1.**
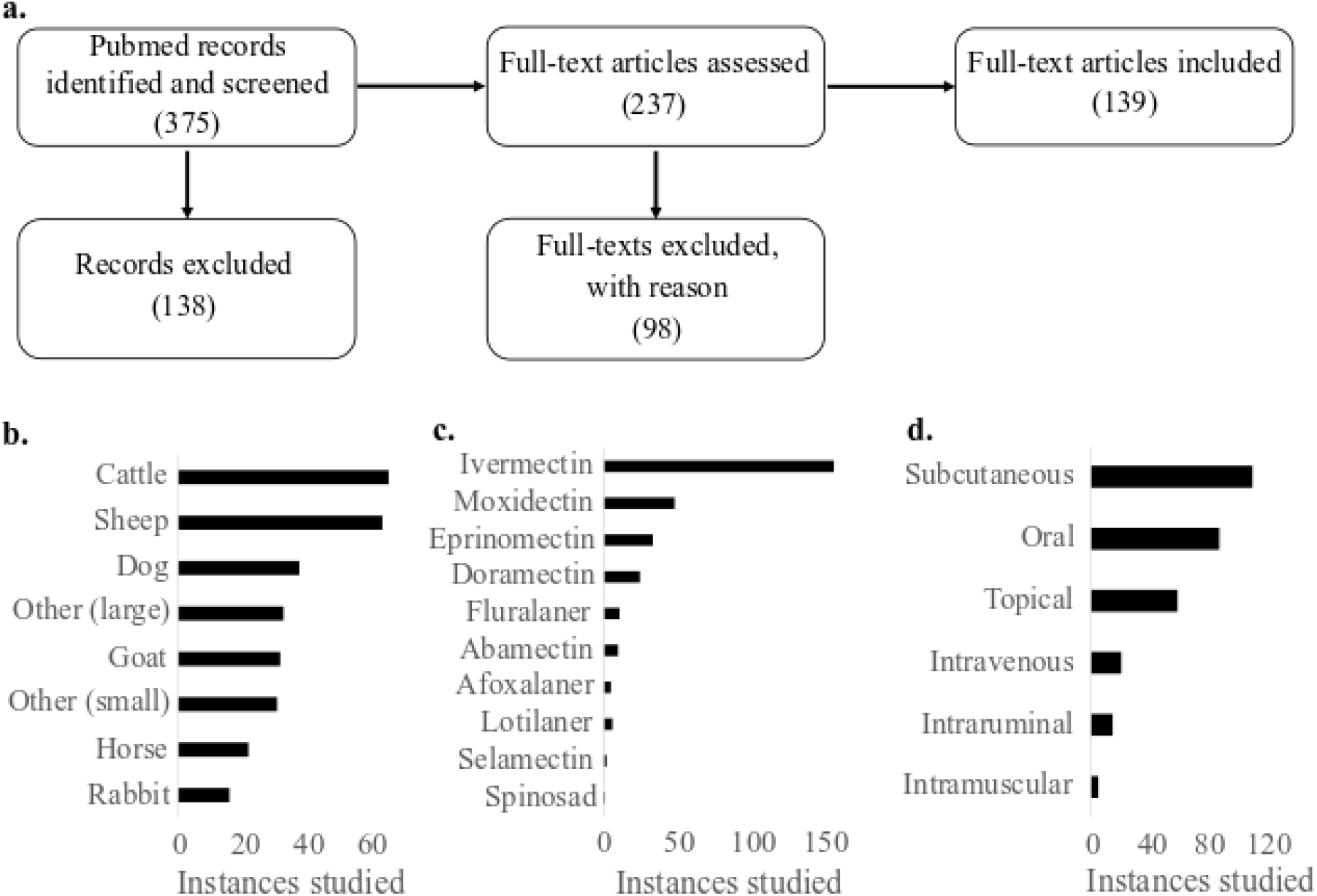
Identification of existing systemic insecticides’ applications. **(a)** A review of PubMed identified relevant studies of existing systemic insecticides. **(b-d)** The included studies covered a range of different hosts, systemic insecticides, and routes of administration.

### II. Data analysis

To evaluate the different treatment scenarios (host, route of administration, and drugs grouped by class), weighted three-way ANOVAs were conducted for each pharmacokinetic parameter. Half-life of elimination and C_max_ were the only parameters with significant interactions (p < 0.05) with host, route, and drug. The significant effectors of half-life were drug class (p = 0.016), route of admission (p = 0.007), and interactions between host and route (p < 0.001) and drug and route (p = 0.007). Regardless of route of administration, the order of drugs from shortest to longest half-lives were avermectins < milbemycins < spinosyns < isoxazolines. The median half-life for avermectins and milbemycins was < 10 days for all routes of administration, whereas the isoxazolines half-lives were > 10 days (Fig. 2a, **Fig. S3**). Comparing median half-lives for a given drug class across hosts shows some host-dependency. For instance, the milbemycin had a longer median half-life in dogs (19.4 days) than in other hosts (< 10 days) and avermectins have a longer median half-life in cattle than in other hosts (Fig. 2b, **Fig. S4**). When comparing routes of administration for different hosts, topically applied drugs typically achieved longer half-lives than orally applied ones (Fig. 2c, **Fig. S5**).

**Fig. 2.**
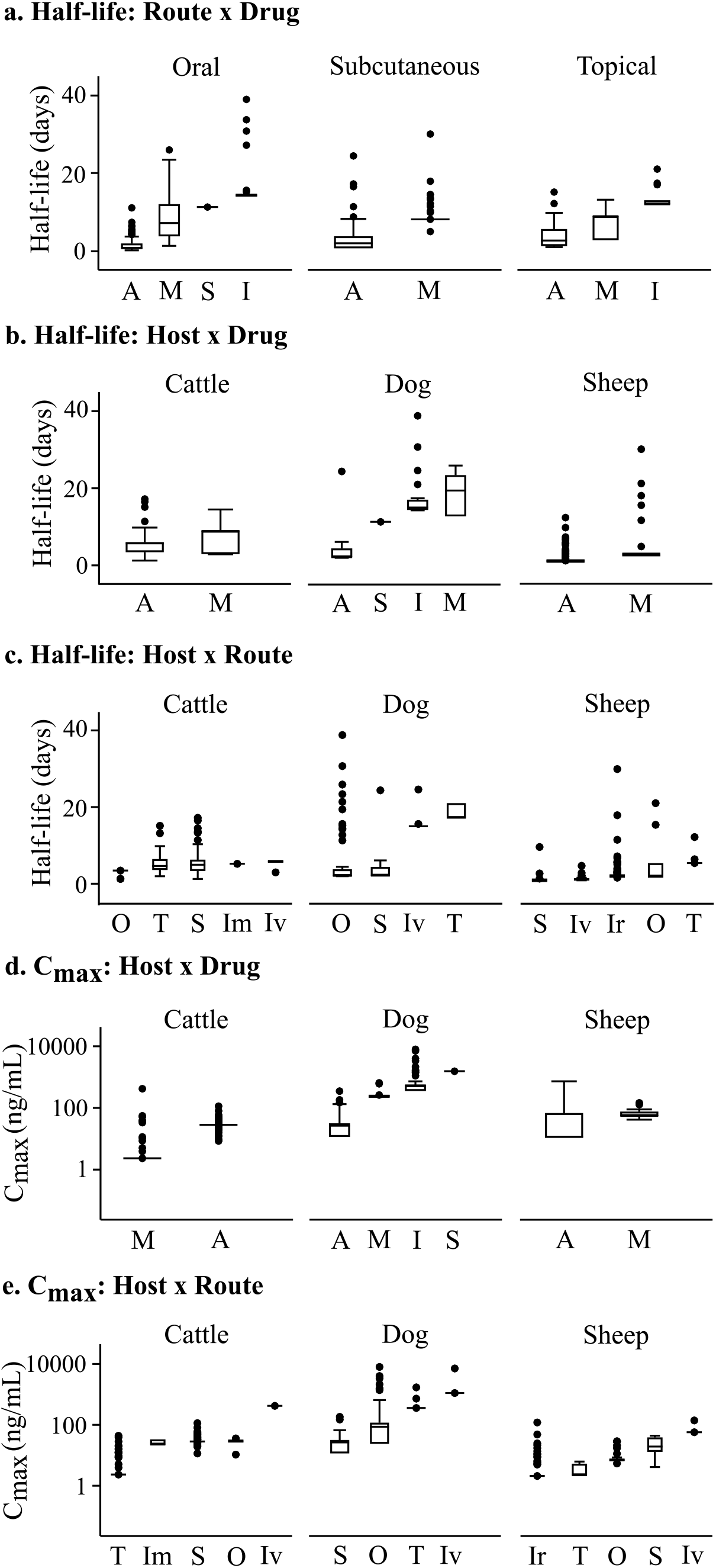
Half-life and C_max_ are dependent on interactions between drug class, host, and route of administration. **(a)** Half-life is affected by the interaction between route of administration and drug class. **(b)** The three most studied hosts show the effect of host and drug class on drug half-life. **(c)** The interaction between drug application route and host affects the drug half-life. (**d**) C_max_ is affected by the interaction between host and drug. (**e**) The interaction between route of administration and host also impacts C_max_. Abbreviations: Drug classes: A= Avermectin, I = Isoxazoline, M=Milbemycin, S= Spinosyn; Routes: Im = Intramuscular, Ir = Intraruminal, Iv = Intravenous, O = Oral, S = Subcutaneous, T = Topical.

The significant factors for C_max_ were drug (p = 0.03) and interactions between host and drug (p = 0.002) and host and route (p < 0.001). The order of drugs from lowest to highest C_max_ was different from that of half-lives: milbemycins < avermectins < isoxazolines < spinosyns (**Fig. S6**). Cattle reported the lowest median C_max_ for milbemycins, whereas dogs and sheep had the lowest C_max_ for avermectins (Fig. 2d, **Fig. S7**). There was also a dependency of C_max_ on host and route (Fig. 2e, **Fig. S8**). Although the intravenous route resulted in the highest C_max_ for different hosts, due to the drug being directly delivered into the bloodstream, the order of resulting C_max_ for other routes varied based on host.

The spread in half-lives and C_max_s seen for a given drug-host-route combination can be attributed to several host factors that may affect some drugs’ absorption and, consequently, the plasma concentration. These factors include age, gender, breed, diet, presence of parasite infection, pregnancy, lactation, and whether or not topically treated hosts are restricted from self-licking (34–41). Understanding how these host conditions affect basic pharmacokinetics is critical for designing optimal treatment strategies that can account for these natural variations.

### III. Basic dynamics of malaria transmission and control methods

The model captures the temporal dynamics of malaria transmission in a human population being exposed to mosquito bites. Upon the introduction of the LLIN, a general decline in malaria prevalence (the sum of symptomatic and asymptomatic individuals) is observed, followed by a steady increase as the insecticide in the net degrades and fewer mosquitoes are killed by LLIN exposure (Fig. 3a). The addition of treating a blood host with systemic insecticide can further reduce malaria prevalence when applied frequently enough.

**Fig. 3.**
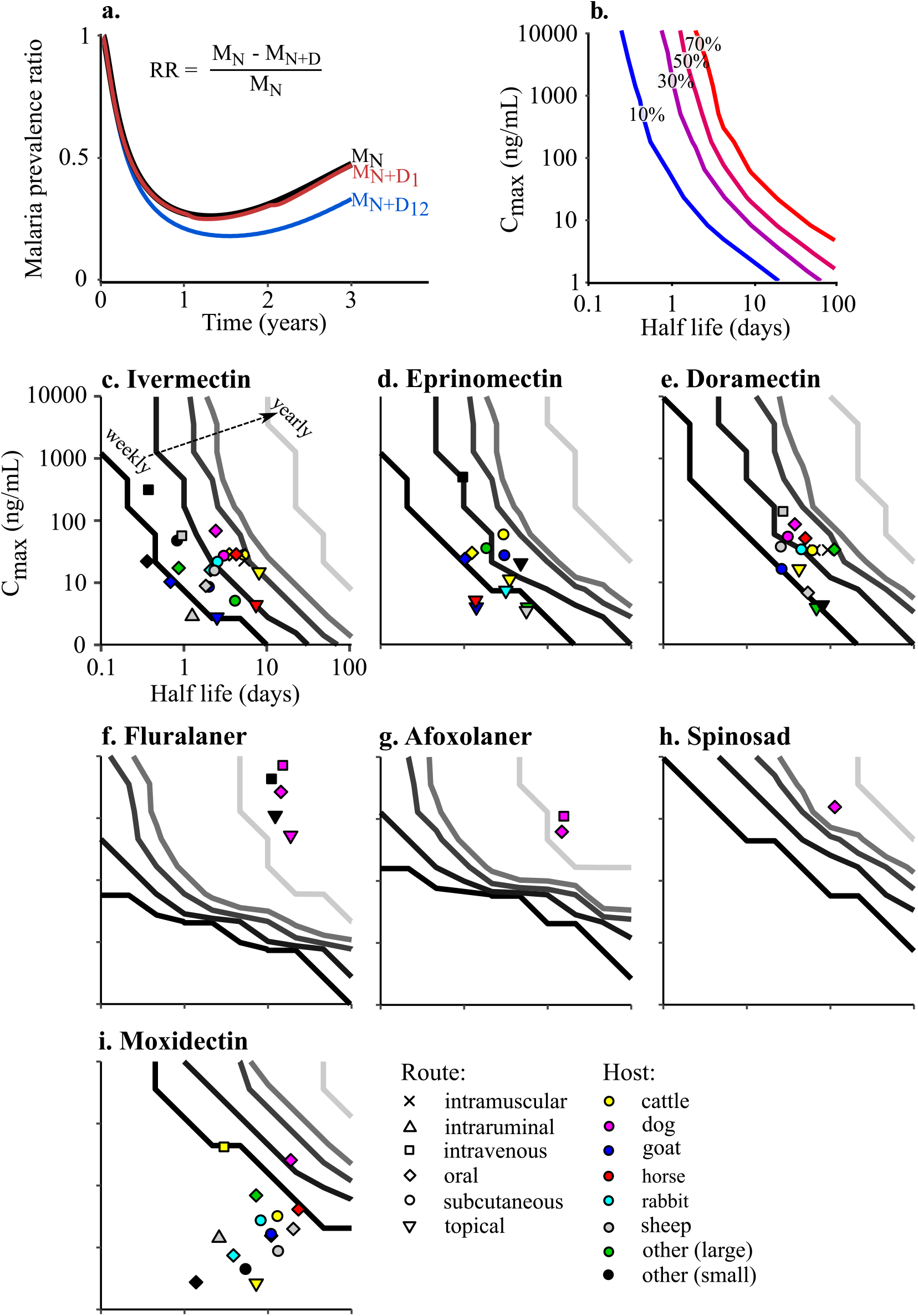
Modelling malaria transmission and control methods. **(a)** The malaria prevalence ratio is compared for a population using LLINs alone (M_N_, black), LLINs with livestock treated yearly with systemic insecticide (M_N+D1_, red), and LLINs with livestock treated monthly with systemic insecticide (M_N+D12_, blue). **(b)** For a strategy using LLINs and livestock treated at a set dosing frequency (here, monthly), the relative reduction in malaria prevalence can be calculated for insecticides of various half-lives and C_max_s. **(c-i)** The dosing frequency necessary to achieve a 10% relative reduction in malaria prevalence can be calculated for insecticides with different pharmacokinetic properties. Overlaying pharmacokinetic values gathered from the review predicts the minimum dosing frequency of existing systemic insecticides in certain host-route scenarios. Contour definitions from left to right: weekly, monthly, quarter-annually, bi-annually, annually. Here, we assume indiscriminate biting behaviour (p_h_ = 0.5) and a mesoendemic environment (m=10).

### IV. Dosing strategy design and evaluation

To quantify the efficacy of different dosing strategies, the relative reduction (RR) in malaria prevalence after three years of using LLINs and systemic insecticide treated livestock was compared to that of using LLINs alone. For a set dosing frequency, the RR increased as a function of half-life and C_max_ (Fig. 3b).

The minimum dosing frequency was calculated to achieve a target RR (here, 10%) for a range of different drugs distinguished by half-life, C_max_, *F*_50_, and 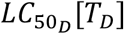 (Fig. 3c-i). The drugs with the longest half-lives and highest C_max_s needed to be dosed the least often to maintain a sufficiently high concentration to remain lethal to feeding mosquitoes. Given the same half-life and C_max_, drugs with higher *F*_50_ and 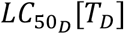 needed to be dosed more frequently to compensate for the decreased efficacy of drug on mosquito fecundity or lethality.

Overlaying the data gathered in the review for drugs with reported *F*_50_s and 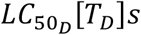 for *Anopheles gambiae sensu lato* shows the frequency at which these existing drugs would need to be applied to achieve the target 10% relative reduction in malaria prevalence. The avermectins, represented by ivermectin, eprinomectin, and doramectin, have relatively low *F*_50_s and 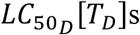, suggesting that relatively low concentrations of drug in the bloodstream would affect the fecundity and death rates of feeding mosquitoes. However, this impact is limited by these drugs’ relatively short half-lives, ranging from 0.4 to 11.1 days (Table 1). Depending on the host and route of administration, regimens with dosing frequencies ranging from weekly to quarter-annually would be required to achieve the target 10% relative reduction in malaria prevalence. Although ivermectin and eprinomectin have a similar 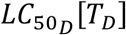, ivermectin has a stronger effect on fecundity (**Fig. S2**). Consequently, ivermectin would require less frequent dosing than eprinomectin to achieve the same target reduction, given scenarios with the same C_max_ and half-life. Doramectin has a lower impact on fecundity and death rates, with higher *F*_50_ and 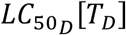 than ivermectin and eprinomectin.

**Table 1.** Range of pharmacokinetic and pharmacodynacmic parameters collected from the literature review.

|  | C<sub>max</sub> (ng/mL) | Half-life (day<sup>-1</sup>) | F<sub>50</sub> (ng/mL) | LC<sub>50</sub>[T<sub>D</sub>] (ng/mL) |
| --- | --- | --- | --- | --- |
| Macrocyclic lactones |  |  |  |  |
| Doramectin | 3.8 : 139.8 | 2.5 : 11.1 | 9.2 | 30.6 |
| Eprinomectin | 3.4 : 503.0 | 1.0 : 5.7 | 1.0 | 7.6 |
| Ivermectin | 2.7 : 316.0 | 0.4 : 7.8 | 4.1 | 7.3 |
| Moxidectin | 2.6 : 420.0 | 1.4 : 23.1 | 478.4 | 1178.0 |
| Isoxazolines |  |  |  |  |
| Afoxolaner | 621.9 : 1107.0 | 14.8 : 15.6 | 0 * | 66.8 |
| Fluralaner | 513.3 : 7109.0 | 11.0 : 18.6 | 0 * | 21.2 |
| Spinosyns |  |  |  |  |
| Spinosad | 1550.0 | 11.3 | 0 * | 461.0 |

\* F<sub>50</sub> remains to be measured for *Anopheles*

Although fluralaner and afoxolaner (both isoxazolines) have higher 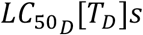 than ivermectin, they are predicted to achieve 10% RR with yearly dosing, due to their longer half-life and higher C_max_. Similarly, spinosad has a high 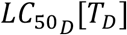, a relatively long half-life of 11.3 days, and a much higher C_max_ of 1550.0 ng/mL, and could achieve a 10% RR when dosed biannually. These results are conservative, as the effect of fluralaner, afoxolaner, and spinosad on fecundity are assumed to be zero until this effect has been characterised in mosquitoes.

Despite some of the studies evaluating moxidectin reported the longest half-lives and highest C_max_s, its high *F*_50_ and 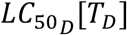 means that it would have to be dosed more frequently (> weekly) or at higher doses to provide an effective complement to LLINs.

### V. Application to different scenarios

This method for evaluating dosing strategies for systemic insecticides can be used to evaluate differences in baseline malaria prevalence in a community, mosquito feeding behaviour, and coverage scenarios. For each scenario, a prediction was made for the dosing frequency necessary for scenarios in which livestock, humans, or both are treated with a systemic insecticide similar to ivermectin (Table 1). Unless otherwise mentioned, it was assumed that mosquitoes were indiscriminate feeders, resistant to permethrin, and malaria was mesoendemic.

#### i. Malaria endemicity

Control methods for varying levels of malaria endemicity were explored by simulating mesoendemic and hyperendemic environments (baseline prevalence between 11% and 50%, or >50%, respectively) (Fig. 4a-c) (42). For all levels of endemecity, treating only livestock or only humans resulted in the same amount of relative reduction for a given half-life and C_max_ because the mosquitoes were simulated as indiscriminate feeders. Treating both livestock and humans had a compounding effect that resulted in the greatest reduction in malaria prevalence and a down-shift in dosing frequencies for a set half-life and C_max_. With increasing malaria prevalence, a drug with the same half-life and C_max_ would need to be dosed more frequently to achieve the same relative reduction in malaria. In a low level mesoendemic environment (malaria prevalence = 25%), LLINs alone play a significant role in reducing transmission; however, additional treatment of livestock and humans could further reduce prevalence and theoretically break transmission. In a high level mesoendemic environment (malaria prevalence = 50%), LLINs alone are not as effective and the additional treatment of livestock and humans could significantly reduce malaria transmission. When malaria is hyperendemic (prevalence = 65%), the addition of systemic insecticide treatment would reduce malaria prevalence relative to LLINs alone; however, to bring malaria transmission under control, longer, sustained treatment and/or the use of drugs with longer half-lives and higher C_max_ would be necessary.

**Fig. 4.**
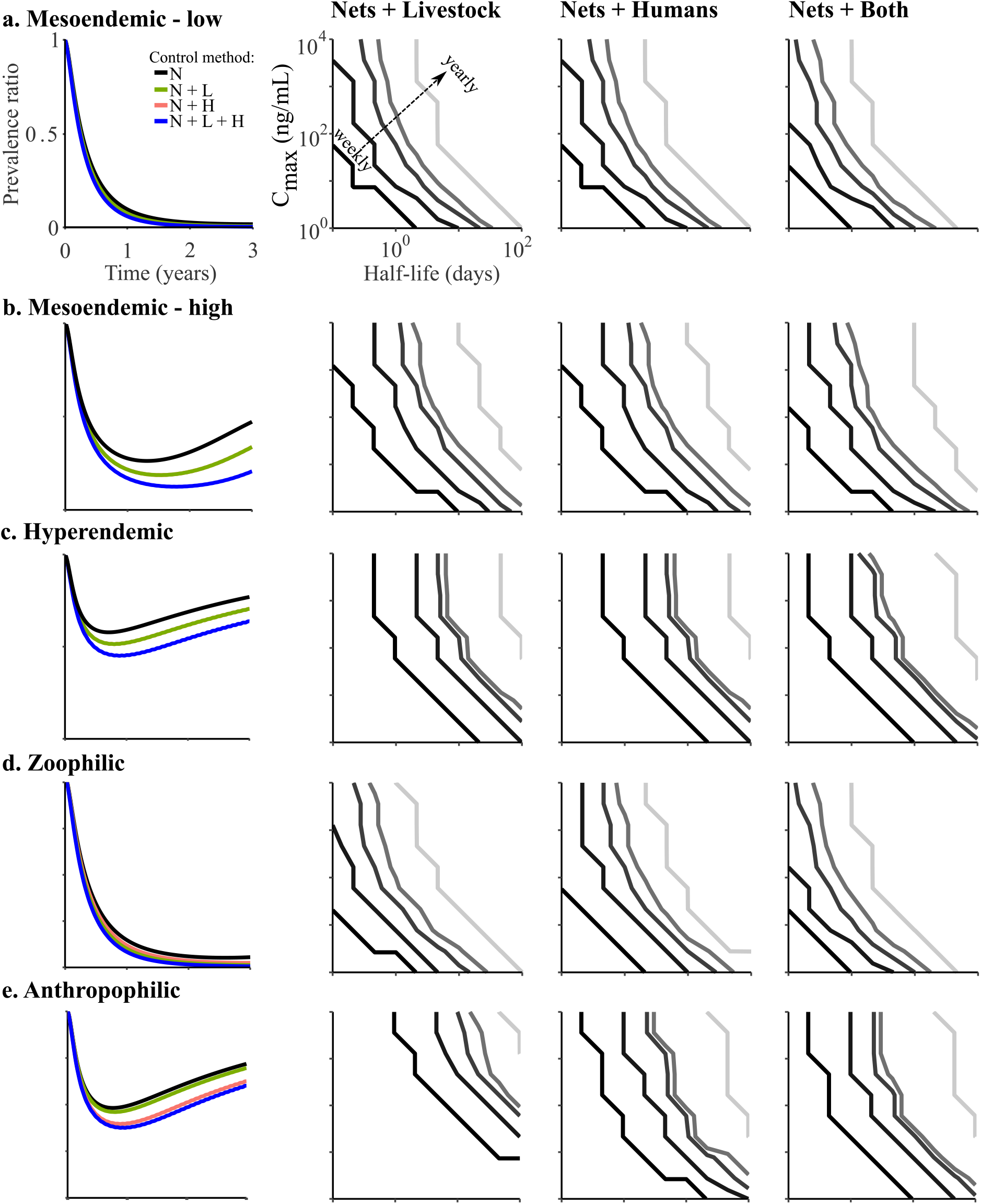
Effect of different coverage strategies for scenarios with different malaria prevalence classes and mosquito biting behaviours. The first column compares temporal dynamics of malaria prevalence for the different scenarios: LLINs alone (N), LLINs and livestock treatment (N+L), LLINs and human treatment (N+H), LLINs with both hosts treated (N + both). The three right columns are the predictions for dosing frequency necessary to reduce malaria prevalence by 10% for each scenario. **(a-c)** With increasing malaria presence, the degree to which control methods can reduce malaria prevalence decreases and the frequency of insecticide reapplication increases (mesoendemic-low: m = 5; mesoendemic-high: m = 10; hyperendemic: m=20). Here, an indiscriminate biting behaviour is assumed in mosquitoes (p_H_ = 0.5), thus N+L and N+H have same outcome. (**d-e**) When mosquitoes are zoophilic (p_H_ = 0.35), systemic insecticides do not need to be dosed in livestock or humans as frequently as in other scenarios, due to lower rates of mosquitoes biting humans. Controlling anthropophilic mosquitoes (p_H_ = 0.8) requires an increase in dosing frequency due to the high rate of human bites. Here, m = 10. Contour definitions from left to right: weekly, monthly, quarter-annually, bi-annually, annually.

#### ii. Mosquito feeding behaviour

Choosing the correct control method for a given community also relies on the bloodmeal preference of the mosquito population (Fig. 4d-e) (43). When mosquitoes are zoophilic, malaria transmission can largely be controlled by LLINs because the mosquitoes do not target humans as frequently. Treating livestock with systemic insecticides would be more effective than treating humans, requiring less frequent dosing for a drug with a given half-life and C_max_. Malaria in regions with anthropophilic mosquitoes was reduced the most with the treatment of both livestock and humans, with most of the reduction due to the treatment of humans. Dosing frequencies were increased, given the need to maintain high enough lethal systemic insecticide concentrations to affect a greater number of mosquitoes targeting humans for bloodmeals.

## Discussion

To date, ivermectin has been the main systemic insecticide considered for its mosquitocidal properties and the only one marketed for human use (44–46). However, there are a range of additional avermectins and different drug classes, such as milbemycins, spinosyns, and isoxazolines, that should be screened as potential candidates for future mosquito control methods in livestock and/or humans. Here, through the combination of a review and a malaria transmission model, it was demonstrated how existing systemic insecticides can be applied to reduce malaria transmission in a range of scenarios. To make a 10% reduction in malaria prevalence beyond what is already achieved by LLINs, some of the identified drugs need to be applied at frequencies much higher than the current MDA’s annual or bi-annual dosing regimen (19). For instance, it was estimated that many scenarios using moxidectin would need to be dosed at an unrealistic frequency, faster than once a week, to reduce malaria transmission. This supports previous studies that show moxidectin has little impact on mosquitoes (28) and would not be an effective choice for a treatment strategy. Further investigation of these drugs’ safety in hosts and effect on mosquitoes is necessary.

To design treatments with an operationally realistic dosing frequency (i.e. once or twice a year), drugs that naturally attain higher blood concentrations and have longer half-lives (i.e. isoxazolines and spinosyns) could be selected or alternative ways to achieve higher concentrations of drug for longer periods of time could be applied. Recently, a study demonstrated the tolerability of high doses of ivermectin in humans and the resulting increase in time that the blood was lethal to mosquitoes (47). An alternative to increasing the concentration of the delivered dose, a number of adjuvants have been reported to improve the absorption or extend the half-life of systemic insecticides, such as lipids for lipophilic drugs (48, 49) or efflux pump inhibitors that remove xenobiotics (50–54), respectively. A third promising solution is the development of sustained-release devices, which have been shown to maintain a target concentration for 280 days in livestock (55).

In addition to designing a dosing regimen that is effective at reducing the transmission of a vector-borne disease, the secondary effects should also be considered:

### Emergence of resistance

As the frequency of insecticide administration is increased to break the malaria transmission cycle, it is imperative to consider how these new treatment regimens will lead to the emergence of resistance in mosquitoes as well as other parasites. To gain some insight, consider how mosquitoes have already developed resistance to the insecticides used in LLINs and IRS (3). The two main mechanisms of insecticide resistance observed so far in mosquitoes can be categorized as metabolic or target site mutations (56). Studying how other arthropods have formed resistance to these systemic insecticides can also shed light on the path of resistance formation in mosquitoes. For instance, macrocyclic lactone resistance has been shown to arise in the cattle tick, fruit fly, and body lice due to increased expression of ATP-binding cassette transporter, P-glycoprotein, and P450 genes (57–59). With strategic use of systemic insecticides that target different mechanisms, monotherapy could be avoided and the development of resistance in mosquitoes could be delayed.

Selecting for resistant off-target parasites is also a concern. One study reported ivermectin resistant *Rhipicephalus microplus* found on 50% of cattle being treated regularly with ivermectin for gastrointestinal nematodes (60). Similarly, repeated treatment of ivermectin for onchocerciasis in humans has selected for resistant *Onchocerca volvulus* in Ghana (61). As these systemic insecticides are commonly used to control other parasite infections in humans and livestock, it is critical that dosing regimens and outcome surveillance addresses the formation of resistance in off-target parasites.

### Presence in food products

Considering how different systemic drugs are secreted from a host’s system is also a critical part of evaluating a new treatment program. For instance, lactating hosts treated with a single-dose of eprinomectin or ivermectin produced milk with detectable drug levels for weeks, thus exposing the nursing young (62, 63). Due to the unknown effects of the drug on a newborn, nursing human mothers in the first week after delivery have been excluded from MDAs of ivermectin (64).

Additionally, systemic insecticides from treated hosts becomes incorporated into dairy and meat products (65). Regulations have been established for levels of acceptable residues of a few drugs, such as eprinomectin; yet, most other drugs have not been licensed for use in dairy animals and thus do not have an acceptable limit for drug concentration found in milk (66). It was surmised that the extent of drug excretion and residence time in milk depends on a drug’s lipophilicity and route of administration (67, 68). However, more studies are needed to characterise the different drugs’ excretion in milk and, subsequently, establish safety limits for suckling young or consumable items.

## Conclusions

Studies have demonstrated that MDAs of systemic insecticides can significantly decrease mosquito population numbers temporarily. To design effective, long-term vector control strategies with systemic insecticides, their pharmacokinetics and pharmacodynamics need to be understood. Here, multiple systemic insecticides with different mechanisms of action and PK/PD characteristics that could be used in MDAs have been highlighted. The simulations provide a foundation from which to further characterise how wild mosquitoes respond to systemic insecticides. Given the history of mosquitoes forming resistance to the insecticides in LLINs and IRS, having a variety of systemic insecticide strategies that target different mechanisms could help reduce the rate at which mosquito resistance arises to these new methods.

## Acknowledgements

HM was funded by the Whitaker International Program. LFK is supported by an Australian National Health and Medical Research Council Fellowship (APP1158469).

