## Supplementary Information for "Optimizing systemic insecticide use to improve malaria control"

### Supplementary Text: Model development and calculations

#### 1. Model development

Humans

$$\frac{dS_h}{dt} = r(E_h + I_h + A_h) + q_2 R_h - m b_h p_h S_h I_m \quad (1)$$

$$\frac{dE_h}{dt} = m b_h p_h S_h I_m - (\xi_h + r) E_h \quad (2)$$

$$\frac{dI_h}{dt} = \xi_h E_h - (q_1 + r) I_h \quad (3)$$

$$\frac{dR_h}{dt} = q_1(I_h + A_h) - (\theta m b_h p_h I_m + q_2) R_h \quad (4)$$

$$\frac{dA_h}{dt} = \theta m b_h p_h R_h I_m - (q_1 + r) A_h \quad (5)$$

$$b_h = a\, b\, (1 - C_N \frac{N^{H_n}}{N^{H_n} + {LC_{50}}_N [T_N]^{H_n}}) \quad (6)$$

Mosquitoes

$$\frac{dO}{dt} = \beta_{mc} V - d_o O - \mu_O \left(1 + \frac{O+L}{K}\right) O \quad (7)$$

$$\frac{dL}{dt} = d_O O - d_L L - \mu_L \left(1 + \gamma \frac{O+L}{K}\right) L \quad (8)$$

$$\frac{dP}{dt} = d_L L - d_P P - \mu_P P \quad (9)$$

$$\frac{dS_m}{dt} = \frac{1}{2} d_P P - b_m p_h (I_h + \sigma A_h) S_m - \mu_{mc} S_m \quad (10)$$

$$\frac{dE_m}{dt} = b_m p_h (I_h + \sigma A_h) S_m - (\xi_m + \mu_{mc}) E_m \quad (11)$$

$$\frac{dI_m}{dt} = \xi_m E_m - \mu_{mc} I_m \quad (12)$$

$$V = S_m + E_m + I_m \tag{13}$$

$$b_m = a\,c\,(1 - C_N \frac{N^{H_n}}{N^{H_n} + LC_{50_N}[T_N]^{H_n}}) \tag{14}$$

Insecticides

$$\frac{dD_L}{dt} = -k_{D_L} D_L \tag{15}$$

$$\frac{dD_H}{dt} = -k_{D_H} D_H \tag{16}$$

$$\frac{dN}{dt} = -k_N N \tag{17}$$

$$\beta_{mc} = \beta_m - \frac{1}{3}\left((1 - p_h)C_L \frac{D_L^{H_b}}{D_L^{H_b} + F_{50}{}^{H_b}} + p_h C_H \frac{D_H^{H_b}}{D_H^{H_b} + F_{50}{}^{H_b}}\right)\beta_m \tag{18}$$

$$\mu_{mc} = \mu_m + \frac{1}{3}\left(\begin{array}{c} p_h C_N \frac{N^{H_n}}{N^{H_n} + LC_{50_N}[T_N]^{H_n}} \mu_N + \\ (1 - p_h)C_L \frac{D_L^{H_d}}{D_L^{H_d} + LC_{50_D}[T_D]^{H_d}} \mu_D + \\ p_h C_H \frac{D_H^{H_d}}{D_H^{H_d} + LC_{50_D}[T_D]^{H_d}} \mu_D \end{array}\right) \tag{19}$$

$$\mu_N = \frac{2\log(2)}{T_N} \tag{20}$$

$$\mu_D = \frac{2\log(2)}{T_D} \tag{21}$$

Susceptible people are exposed ($E_h$) to malaria as a function of the proportion of bites that successfully leads to infection ($b_m$), the proportion of bites that are made on humans ($p_h$), the ratio of vectors to bloodmeal sources ($m$), and the number of infected mosquitoes ($I_m$). The transmission potential from vector to human or vice versa ($b_h$ and $b_m$, respectively) is a function of biting frequency ($a$), the proportion of bites that successfully leads to infection in humans or mosquitoes ($b$ and $c$, respectively), the coverage of bed nets ($C_N$) and whether the LLIN's insecticidal concentration is above the lethal dose for 50% ($LC_{50_N}[T_N]$) of the population. Exposed humans can either recover to the susceptible state at rate $r$ or transition to the infected state ($I_h$) at the rate at which the parasite matures ($\xi_h$). Infected humans can recover to the susceptible state or form immunity ($R_h$) at rate $q_1$ that is temporary (waning at rate $q_2$) and partial (as determined by at rate $\theta$). Asymptomatic ($A_h$) humans can recover to a susceptible state or return to the immune state. The net change of human populations was assumed to be negligible for the timeframe observed.

All mosquitoes (V), regardless of their infection status, lay $\beta_{mc}$ eggs per day. Coverage of systematic insecticides in livestock ($D_L$) or humans ($D_H$) ($C_L$ and $C_H$, respectively) will affect the egg natural laying rate ($\beta_m$) if the drug concentration is above the concentration associated with a 50% fecundity reduction ($F_{50}$). Eggs hatch into early larval instars (O) that either develop into late instars (L) at rate $d_O$ or die at a rate $\mu_O$. Late larval instars develop into pupae (P) at rate $d_L$ and die at rate $\mu_L$. The death rates of the instar larval stages are dependent on the density of early and late larval instars and the environmental larval carrying capacity (K). To capture the different density dependences, the late larval instars' death rate is modified by $\gamma$. Pupae develop into sensitive mosquitoes ($S_m$, 50% of pupae are assumed to be female) at a rate $d_P$ and die at rate $\mu_P$. Sensitive mosquitoes are exposed ($E_m$) to malaria as a function of biting frequency, the proportion of bites that successfully lead to infection ($c$), the proportion of bites on humans, the ratio of vectors to bloodmeal sources, and the number of infected humans ($I_h$ and $A_h$). Biting an asymptomatically infected human has a lower likelihood of leading to infection and is adjusted by factor $\sigma$. Mosquitoes become infectious at the rate of parasite development ($\xi_m$). Mosquitoes die at a natural rate of $\mu_m$, but the coverage of LLINs and systematic insecticides in livestock ($D_L$) or humans ($D_H$) ($C_N$, $C_L$, $C_H$, respectively) will affect this if the drug concentrations are above the lethal dose for killing 50% of the population in a set period of time ($LC_{50_N}[T_N]$ and $LC_{50_D}[T_D]$). When the mosquitoes are exposed to sufficiently high doses of drug, the maximum death rates due to LLINs ($\mu_N$) and insecticides ($\mu_D$) are approached. $N_{death}$ and $D_{death}$ are dependent on the lethal concentration for killing 50% of exposed mosquitoes ($LC_{50_N}[T_N]$ and $LC_{50_D}[T_D]$), the maximum death rate observed due to the insecticide in LLINs or systemic insecticides ($\mu_N$ and $\mu_D$, respectively), and the time window over which the $LC_{50}$ was determined ($T_N$ and $T_D$). The insecticide in the LLINs and systemic insecticides in the host degrade at rates $k_N$, $k_{D_L}$,and $k_{D_H}$, respectively. Initial conditions of $S_h(0) = 0.96$, $E_h(0) = I_h(0) = R_h(0) = A_h(0) = O(0) = L(0) = P(0) = S_m(0) = E_m(0) = I_m(0) = 0.01$ were used for all simulations.

### 2. Calculating $LC_{50}[T]$ of permethrin

Most studies reporting the $LC_{50}[T]$ of permethrin in bednets use the WHO bioassay for determining resistance in mosquitoes. The bioassay protocol calls for 1 hour of exposure time, which is much longer than mosquitoes normally rest on a bed net (3-15 minutes)[1217]. As such, we used a study that observed the number of mosquitoes killed when exposed to a LLIN for 3 minutes via a cone test [18]. Based on the number of mosquitoes killed in 24 hours after this exposure, the death rate ($\mu$) and concentration ($LC_{50}[T]$) associated with killing 50% of an exposed mosquito population (M) in 24 hours post-exposure was derived. The death rate is dependent on drug (D) concentration.

1. Assume mosquito population follows exponential decay

$$M(t) = M(0)e^{-kt} \text{ where } k = \mu\left(\frac{D}{D+LC_{50}}\right)$$

2. Solve for $\mu$ when $D = LC_{50}$ and $M(t) = \frac{M(0)}{2}$ and $t = 1$ day

$$\mu = \frac{2\ln(2)}{\text{t}} = 1.386/day$$

3. Solve for $LC_{50}$ , assuming $\mu = 1.386/day$ and using parameters from Omondi study: D = 2%, M(t) =0.7, M(0), and t = 1 day

$$LC_{50} = D\left(\frac{\mu t}{-\ln\left(\frac{0.9M(0)}{M(0)}\right)} - 1\right) = 2\left(\frac{1.386}{\ln\left(\frac{1}{0.3}\right)}\right) = 3.426\%$$

The hill coefficient associated with the permethrin concentration and mortality observed was determined by fitting data with a nonlinear model. See **Fig. S1g-h**.

**Table S1**: Parameter values and definitions

| Parameter | Definition | Ref. |
|---|---|---|
| $a$ = 0.2 | Biting frequency [ 0.01 – 0.5 day$^{-1}$] | [1] |
| $b$ = 0.5 | Proportion of bites that produce infection in humans [0.2 – 0.5] | [1] |
| $c$ = 0.5 | Proportion of bites that produce infection in mosquitoes [0.5] | [1] |
| $d_O$ = 0.15 | Rate at which early larval instars mature into late larval instars [days$^{-1}$] | [2] |
| $d_L$ = 0.27 | Rate at which late larval instars mature into pupae [days$^{-1}$] | [2] |
| $d_P$= 1.56 | Rate at which pupae mature into adult mosquitoes [days$^{-1}$] | [2] |
| $m$ = 10 | Ratio of mosquitoes to blood meal sources [0.5 - 40] | [1] |
| $p_H$ = 0:1 | Availability of humans to all available blood hosts | [3,4] |
| $q_1$ = 1/200 | Rate of immunity acquisition [days$^{-1}$] | [5] |
| $q_2$ = 1/1000 | Rate of immunity loss [days$_{-1}$] | [5] |
| $r$ = 0.01 | Rate of recovery [0.005 – 0.05 day$^{-1}$] | [1] |
| $\beta_m$ = 21.19 | Number of eggs a mosquito lays per day | [2] |
| $\theta$ = 0.5 | Level of reduced susceptibility to secondary infection | [5] |
| $\sigma$ = 0.25 | Adjustment factor for asymptomatic infection transmissibility to vector | [5] |
| $\mu_O$ = 0.034 | Death rate of early larval instars at low density [days$^{-1}$] | [2] |
| $\mu_L$= 0.035 | Death rate of late larval instars at low density [days$^{-1}$] | [2] |
| $\gamma$ = 13.25 | Factor to correct for different density dependent death rate of late vs. early larval instars [unitless] | [2] |
| $\mu_P$= 0.25 | Death rate of pupae [days$^{-1}$] | [2] |
| $\mu_m$ = 0.12 | Death rate of mosquitoes [0.05 - 0.5 day$^{-1}$] | [1] |
| $\xi_m$ = 1/10 | Rate of *P. falciparum* maturation, given that latent period for mosquitoes is 5 - 15 days [days$^{-1}$] | [1] |
| $\xi_h$ = 1/21 | Rate of *P. falciparum* maturation, given that latent period for humans 10 - 100 days [days$^{-1}$] | [1] |
| $F_{50}$ = 0:8.5 | Threshold systemic insecticide concentration for reducing fecundity by 50% [ng/mL]:<br>$F_{50}$ = 3.6 ng/mL for ivermectin | [9] |

| | | |
|---|---|---|
| | $F_{50}$ = 1.1 ng/mL for eprinomectin<br>$F_{50}$ = 9.2 ng/mL for doramectin<br>$F_{50}$ = 478.4 ng/mL for moxidectin<br>$F_{50}$ = 0 ng/mL for fluralaner, afoxolaner, and spinosad. A conservative estimate, since these values need to be measured | |
| $LC_{50_D}[T_D]$= 7:1180 | $LC_{50}$ of systemic insecticides for *Anopheles* observed over set time [ng/mL]<br>$LC_{50}[9]$ = 7.4 ng/mL for ivermectin<br>$LC_{50}[9]$ = 7.6 ng/mL for eprinomectin<br>$LC_{50}[1]$ = 21.2 ng/mL for fluralaner<br>$LC_{50}[9]$ = 30.6 ng/mL for doramectin<br>$LC_{50}[1]$ = 66.8 ng/mL for afoxolaner<br>$LC_{50}[5]$ = 461 ng/mL for spinosad<br>$LC_{50}[9]$ = 1178 ng/mL for moxidectin | [9–11] |
| $T_D$= 1 : 9 | Time over which systemic insecticide $LC_{50}$s were observed [days]<br>$T_D$= 1 days for afoxolaner and fluralaner<br>$T_D$= 5 days for spinosyn<br>$T_D$= 9 days for doramectin, eprinomectin, ivermectin, and moxidectin | [9–11] |
| $LC_{50_N}[T_N]$ = 3.43 | $LD_{50}$ for permethrin resistant mosquitoes exposed to Olyset nets observed over set time [%] (0.08% for susceptible). Assuming 30% of mosquitoes are killed in 24 hours following 3-minute exposure to a new Olyset net (2% permethrin). | [12], see next section for calculation |
| $T_N$ = 1 | Time over which $LC_{50}$ for permethrin was observed [days] | [12] |
| $H_b = 0{:}16$ | Hill coefficient for systemic insecticides and fecundity [unitless] | [9] |
| $H_d = 1{:}8.5$ | Hill coefficient for systemic insecticides and death rate [unitless] | [9–11] |
| $H_n$ = 2 | Hill coefficient for resistant mosquitoes and permethrin in LLINs ($H_n$= 4 for susceptible) [unitless] | [12] |
| $C_N$ = 0:0.75 | Coverage by bed nets (0:1) * max efficacy of new LLIN (0.75) | [5] |
| $C_L$= 0:1 | Livestock coverage by systemic insecticides (0:1) * max efficacy of newly applied drug (1) | [5] |
| $C_H$ = 0:1 | Human coverage by systemic insecticides (0:1) * max efficacy of newly applied drug (1) | [5] |
| $k_N = \frac{\ln(2)}{1906}$ | After 7 years, Olyset nets had an average concentration of 6.6 g/kg permethrin[14]. With an initial concentration of 20 g/kg[15], the half life is calculated to be 5.2 years or 1906 days. | [15,16] |
| $k_L; k_H$=0.01:10 | Rate of degradation of insecticide in host based on half lives recorded from literature search [$days^{-1}$] | From systematic review |
| $K$ = 3 | Environmental larval carrying capacity. Ranges from 1:10+ | varied |

| | | |
|---|---|---|
| *Net* = 2 | Concentration of permethrin in new Olyset net [%] (1000 mg/$m^2$ or 20 g/kg) | [15] |
| *Dose* = 1:$10^5$ | Concentration of insecticide at start of treatment based on $C_{max}$ measured from literature search [ng/mL] | From systematic review |
| $period_N$ = 1095 | Recommended time between replacement Olyset nets [days] | [15] |
| $period_D$ = 7:365 | Time between administering new dose of systemic insecticide [days] | varied |

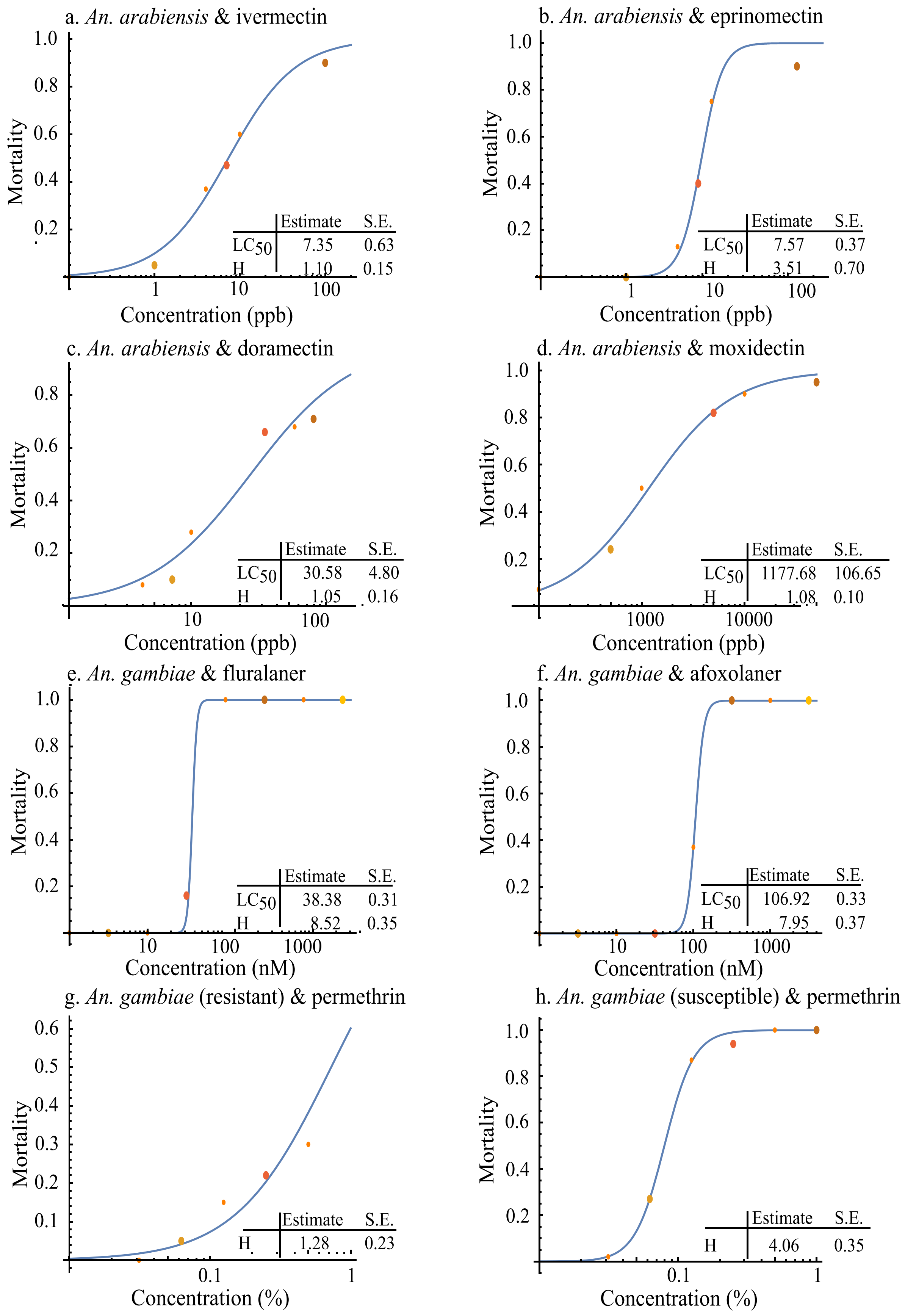

a. An. arabiensis & ivermectin
Mortality
Concentration (ppb)
Estimate S.E.
LC50 7.35 0.63
H 1.10 0.15
b. An. arabiensis & eprinomectin
Mortality
Concentration (ppb)
Estimate S.E.
LC50 7.57 0.37
H 3.51 0.70
c. An. arabiensis & doramectin
Mortality
Concentration (ppb)
Estimate S.E.
LC50 30.58 4.80
H 1.05 0.16
d. An. arabiensis & moxidectin
Mortality
Concentration (ppb)
Estimate S.E.
LC50 1177.68 106.65
H 1.08 0.10
e. An. gambiae & fluralaner
Mortality
Concentration (nM)
Estimate S.E.
LC50 38.38 0.31
H 8.52 0.35
f. An. gambiae & afoxolaner
Mortality
Concentration (nM)
Estimate S.E.
LC50 106.92 0.33
H 7.95 0.37
g. An. gambiae (resistant) & permethrin
Mortality
Concentration (%)
Estimate S.E.
H 1.28 0.23
h. An. gambiae (susceptible) & permethrin
Mortality
Concentration (%)
Estimate S.E.
H 4.06 0.35

**Fig. S1.** Estimating the hill coefficient (H) and $LC_{50}$[T]. **(a-h)** The mortality of mosquitoes for a range of concentrations was plotted for each systemic insecticide, recorded from [9,11,12]. Mathematica was then used to fit a nonlinear model to these data and estimate H and $LC_{50}$[T]. Please note that the $LC_{50}$[T] for permethrin was not calculated based on this data, due to the extended periods of exposure in the protocol. Please see previous section for calculations.

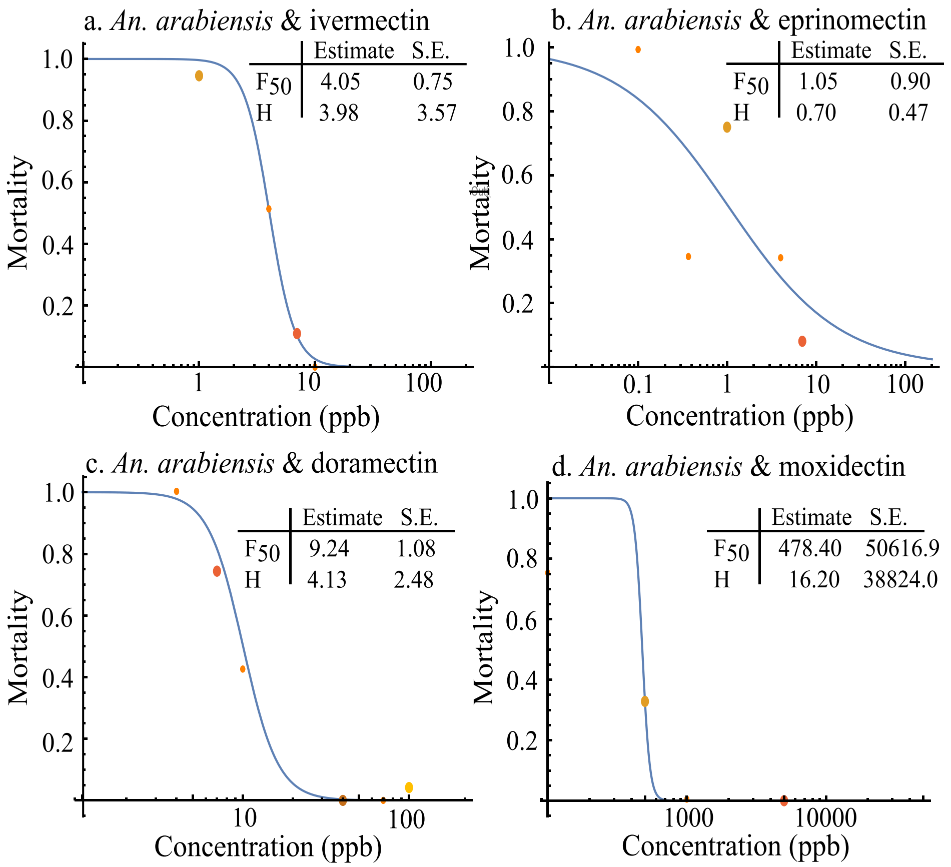


**Fig. S2**. Estimating the hill coefficient (H) and concentration necessary to reduce number of eggs laid per day by 50% ($F_{50}$). **(a-d)** The mortality of mosquitoes for a range of concentrations was plotted for each systemic insecticide, recorded from [9]. A reduction in eggs laid per day has either not been observed or yet to be characterized for permethrin, the isoxazolines, and spinosad. Mathematica was used to fit a nonlinear model to these data and estimate H and $F_{50}$.

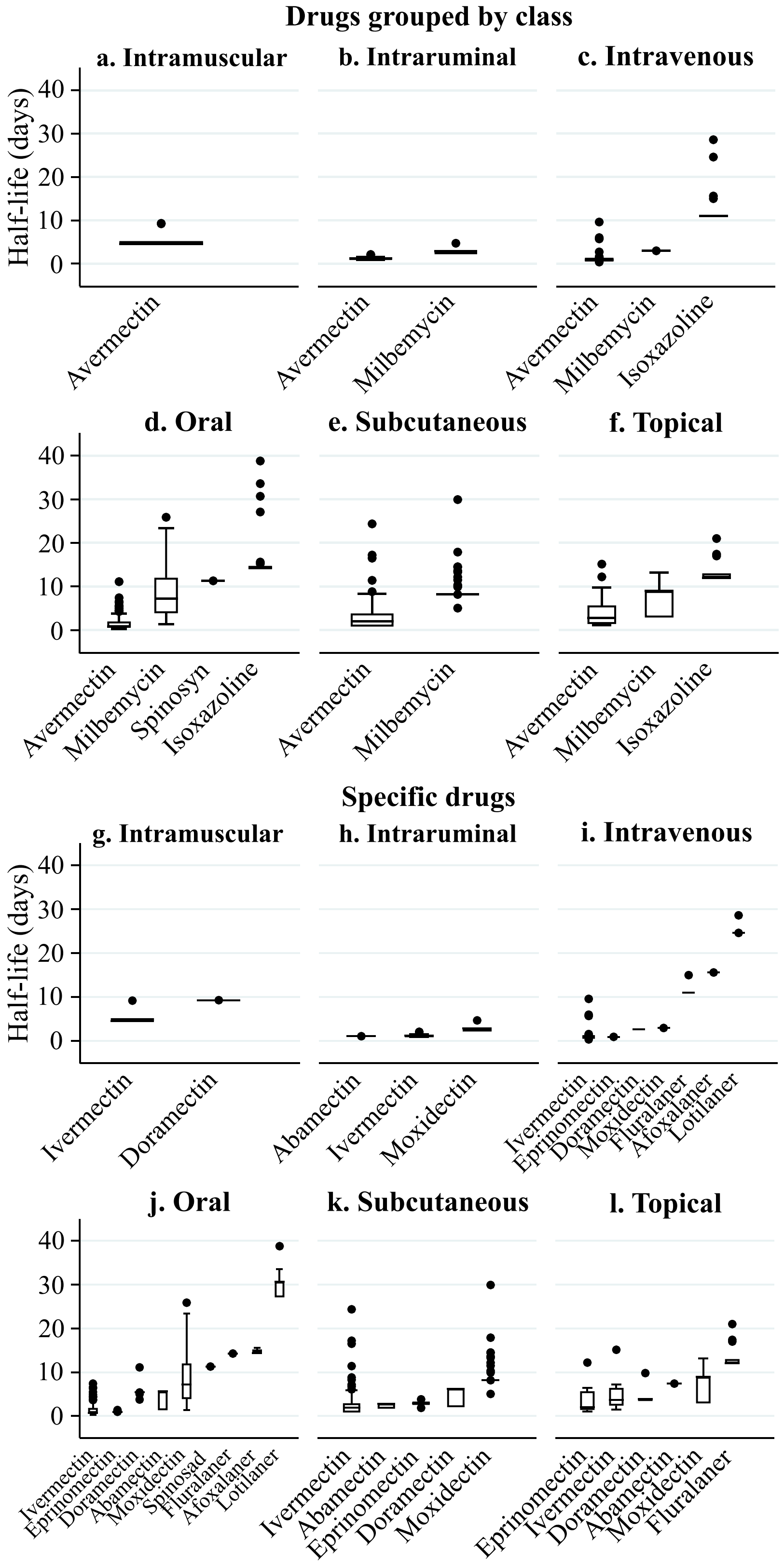


**Fig. S3**. Interactions between route and drug on half-life. (**a-f**) Drugs are grouped into classes; (**g-l**) specific drugs are presented.

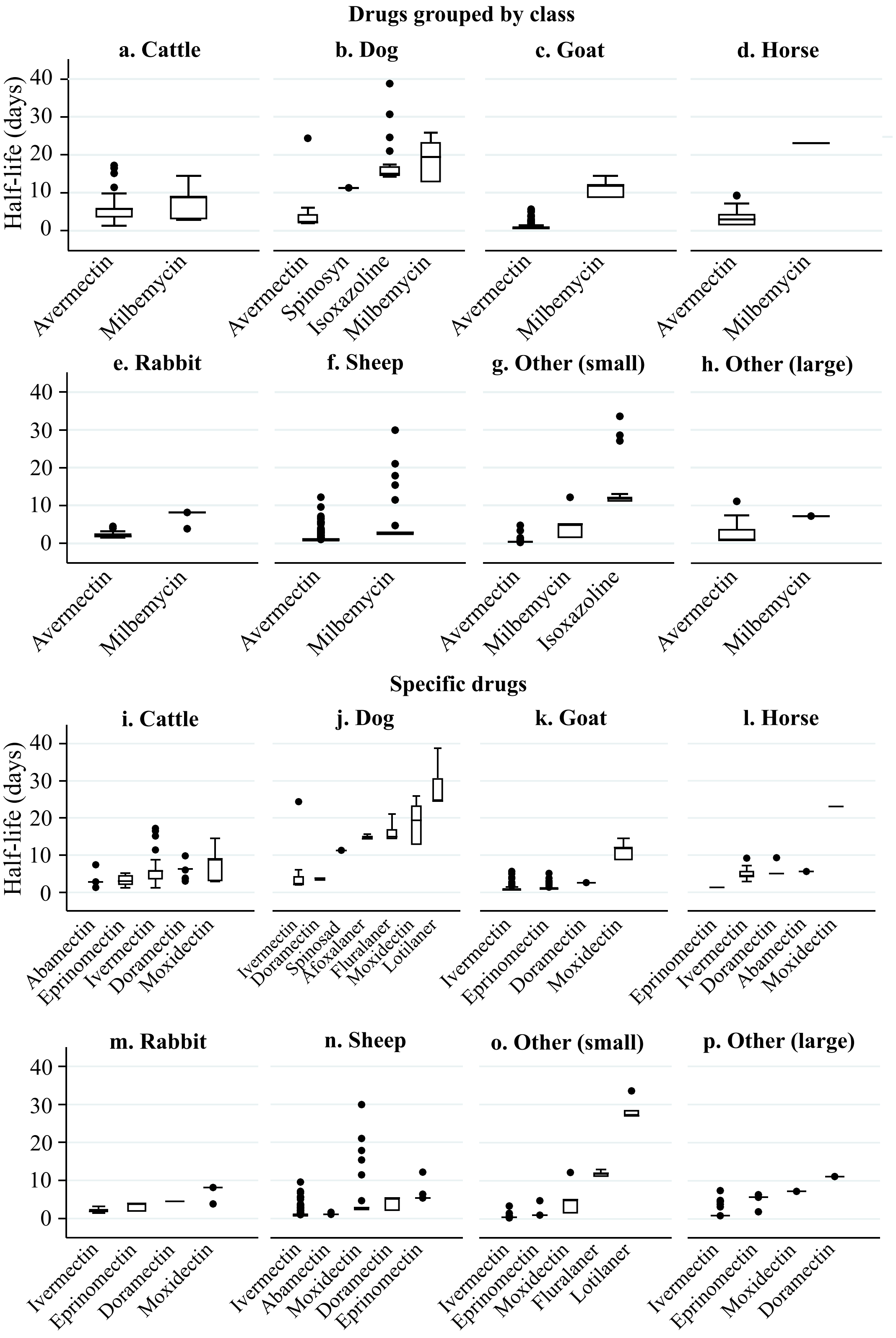


**Fig. S4.** Effect of interactions between host and drug on half-life. (a-h) Drugs are grouped into classes; (i-p) specific drugs are presented.

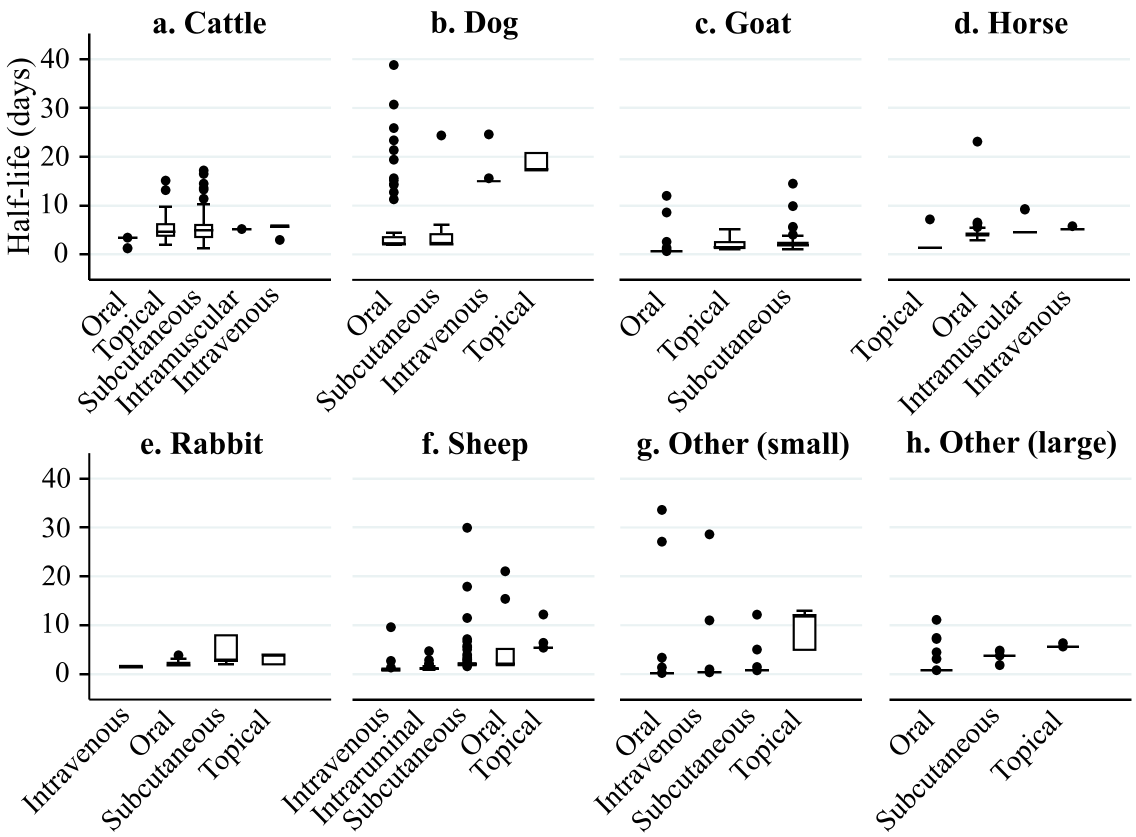


**Fig. S5**. Effect of route and host on half-life.

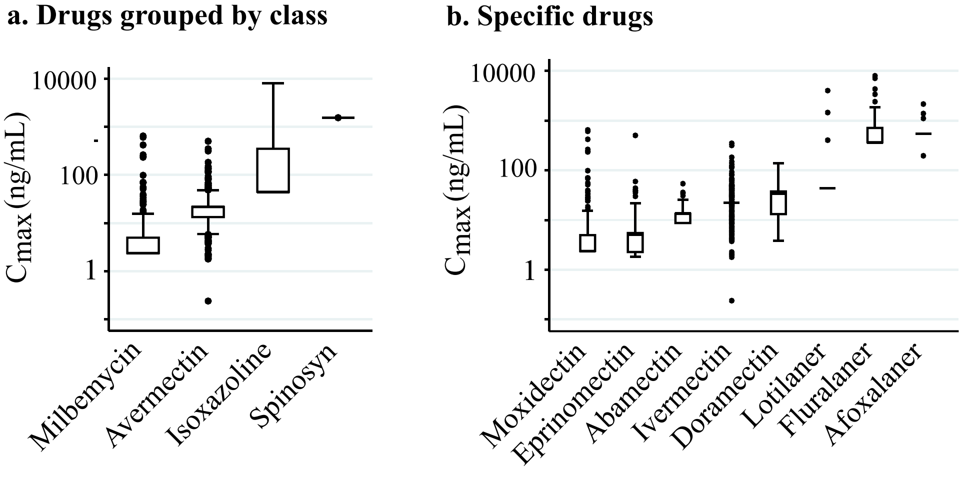


**Fig. S6**. Effect of drug (a. class or b. specific) on $C_{max.}$

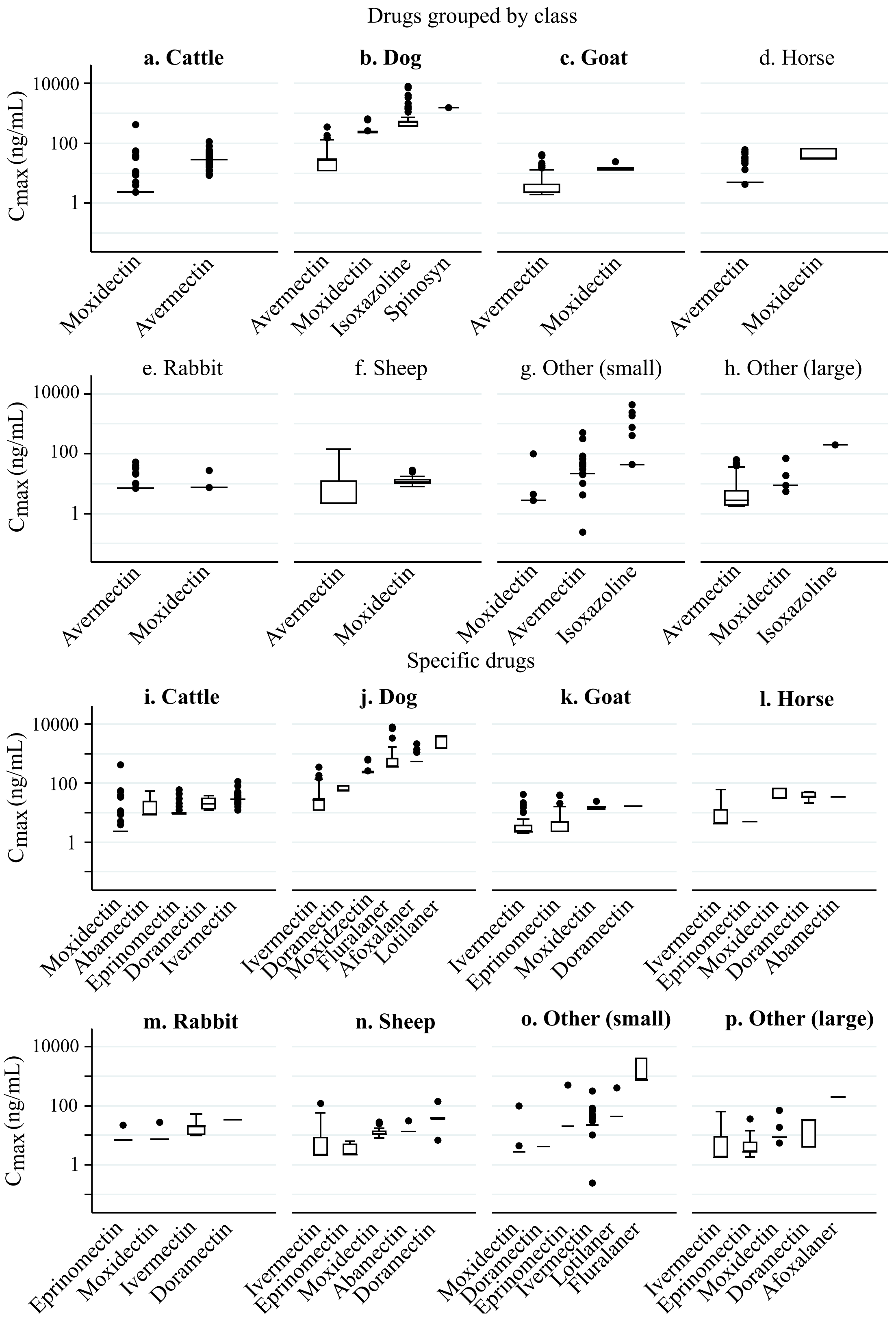


**Fig. S7**. Effect of interactions between host and drug on $C_{max}$. (**a-h**) Drugs are grouped into classes; (**i-p**) specific drugs are presented.

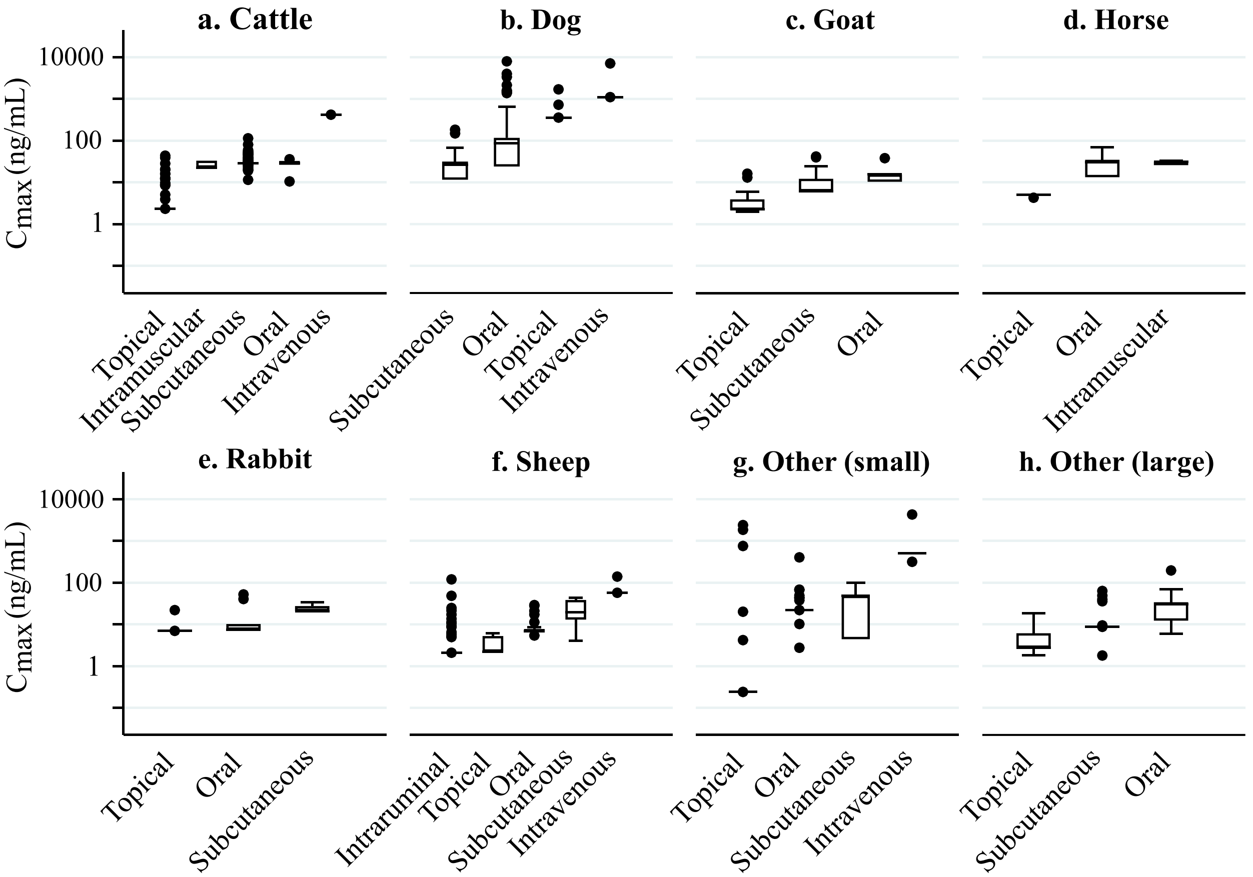


**Fig. S8.** Effect of host and route on $C_{max}$
